# Computational analysis of transition temperatures (Tts) of proteins fused to elastin-like polypeptide (ELP): deep fake evaluation of proteins, linkers, and trailers features

**DOI:** 10.1101/2021.12.29.474407

**Authors:** Mohammad Davoud Ghafari, Iraj Rasooli, Khosro Khajeh, Bahareh Dabirmanesh, Mohammadreza Ghafari, Parviz Owlia

## Abstract

The phase transition temperature (Tt) prediction of the Elastin-like polypeptides (ELPs) is not trivial because it is related to complex sets of variables such as composition, sequence length, hydrophobic characterization, hydrophilic characterization, the sequence order in the fused proteins, linkers and trailer constructs. In this paper, two unique quantitative models are presented for the prediction of the Tt of a family of ELPs that could be fused to different proteins, linkers, and trailers. The lack of need to use multiple software, peptide information, such as PDB file, as well as knowing the second and third structures of proteins are the advantages of this model besides its high accuracy and speed. One of our models could predict the Tt values of the fused ELPs by entering the protein, linker, and trailer features with R2=99%. Also, another model is able to predict the Tt value by entering the fused protein feature with R2=96%. For more reliability, our method is enriched by Artificial Intelligence (AI) to generate similar proteins. In this regard, Generative Adversarial Network (GAN) is our AI method to create fake proteins and similar values. The experimental results show that our strategy for prediction of Tt is reliable in large data.

**Author Summary:** The application of Elastin-like polypeptides (ELPs) as a protein tag is now developed in a variety of biotechnology aspects especially in proteins purification and drug delivery. ELPs application as a protein tag is owed to retain the phase transition behavior when ELPs fused to other proteins, linkers and tags. ELPs undergo the phase transition behavior by changing from soluble phase to insoluble phase above its inverse transition temperature (Tt) within a short time span. This biophysical behavior is usually reversible at the temperature below the Tt. There are few reports for evaluation of the Tt of ELPs types by using the dissimilar equations and algorithms. Our current predictions are the most accurate calculations presented so far by using the protein, linker and trailer effects and the results were evaluated in accordance with the available experimental data. Furthermore, our results also show that our strategy for prediction of Tt is reliable in large data.

## Introduction

ELPs are short artificial repeating peptide motifs classified as biodegradable biopolymers (1, 2). ELPs do not have any adverse immune responses in humans; hence, they can be administered efficiently as a fused tag to biological drugs (3, 4). Also, ELPs are used for enzyme recycling, tissue engineering, and protein purification (5). These polypeptides have shown the minimum immune responses in animal models (6, 7). They undergo phase transition behavior by changing from soluble to insoluble phase above its inverse transition temperature (Tt) within a brief time span. This biophysical behavior is usually reversible at temperatures below the Tt value. The ELP motifs are generally composed of multiple repeats of the pentapeptide sequences (Val-Pro-Gly-Xaa-Gly)n (VPGXG)n. Xaa is a guest residue, while n depicts the number of repeats which is usually between 20-330 residues (8). The guest residue could be any amino acid except proline. It has been demonstrated that proline destroys the inverse phase transition effects (8–11). These repetitive ELP peptides are derived from the native repeated VPGVG motifs of mammalian elastin (12–15).

ELPs can be widely used as a protein tag for protein purification and drug delivery. The application of ELP, as a protein tag, is employed for maintaining the phase transition behavior when ELPs are fused to other proteins, linkers, and tags. Hence, they are characterized by their applications as a result of their simplicity, inexpensiveness, and lack of need for chromatographic purifications of fused proteins (9, 16, 17). This intelligent purification technique is termed Inverse Transition Cycling (ITC) (18). A transition temperature (Tt) is a very narrow temperature range (2–3 °C) used to trigger the phase transition from the soluble to insoluble state. The Tt prediction is not simply obtained because it is related to a complex set of variables, such as the composition, sequence length, hydrophobic characterizations, hydrophilic properties, and the sequence order in the fused proteins. The change in the Tt value can influence the quantity and quality of the ultimate product. For instance, 1) the protein activity is lost or diminished when the Tt value exceeds the required temperature for the activity of a heat-sensitive protein; 2) protein degradation occurs when a high temperature of Tt is required to trigger the phase transition in the process of protein purification; 3) during the protein expression, the phase transition occurs at the growth temperature of bacteria. Therefore, the computational analyses would be significant for researchers to predict such values. Different computational models have previously been reported for the prediction of Tt values in ELPs using dissimilar equations and algorithms. For example, the non-linear models have been reported in two pH-sensitive ELPs by Support vector machine (SVM) and backpropagation neural network (BPNN) algorithms (19). Also, *Chilkoti* et al. quantified the effects of the amino acid sequence and chain length of ELPs (including Ala and/ or Val residues at the guest residue position of pentapeptide sequence) on the Tt values (20).

ELP90 is composed of 90 pentameric ELP repeats and commonly used as an ELP tag. It has Val, Ala, or Gly residues at the fourth position of the pentapeptide sequence. The ratios of these residues are usually equal to 50%, 20%, and 30%, respectively (9, 17, 20–22). ELP90 has wider applications than other ELPs since it provides high yields than many fusion proteins in the T7 expression system in *E. coli* (17). *Chilkoti* et al. indicated that different target proteins, linkers, and tags dramatically altered the Tt value of the fused ELP90 compared with that of the free form of ELP90. They found a linear correlation between the charged amino acids surface index (SIc) of the protein and the values of Tt (23).

To the best of our knowledge, there are no reports on the prediction of the Tt values when different proteins, linkers, and trailers were applied as a quantitative model. Also, no study has been conducted on the possibility of using Weka software for the prediction of Tt. Hence, in this study, two unique quantitative models were introduced to predict the Tt value of the ELP tag fused to different proteins, linkers, and trailers. This model has the following characteristics:

Good ability to calculate the protein, linker, and trailer effects and their impacts on Tt values

No need to use multiple software programs for Tt prediction

No need to know the information about the structure of the protein, such as PDB file, secondary, tertiary, and quaternary structures, to predict the Tt values

It has high accuracy for Tt prediction (i.e., over 96 %)

## Results

### Feature Extraction

The results of ASA and SI calculations are shown for the protein, linker, and trailer features in the Supplementary File. The numerical values of proteins, linkers, trailers, and class attributes were briefly shown in Table 1. The ΔT values shown in the class column were gradually increased from the top to bottom in Table 1. The highest and lowest ΔT values correspond to Barstar and Tendamistat proteins, respectively. The Data Table was added to the Weka software by selecting the pre-process tollbar, open file tab, and the created data table .csv file (data Table .csv file created by the Excell software) (22, 23).

**TABLE 1.**
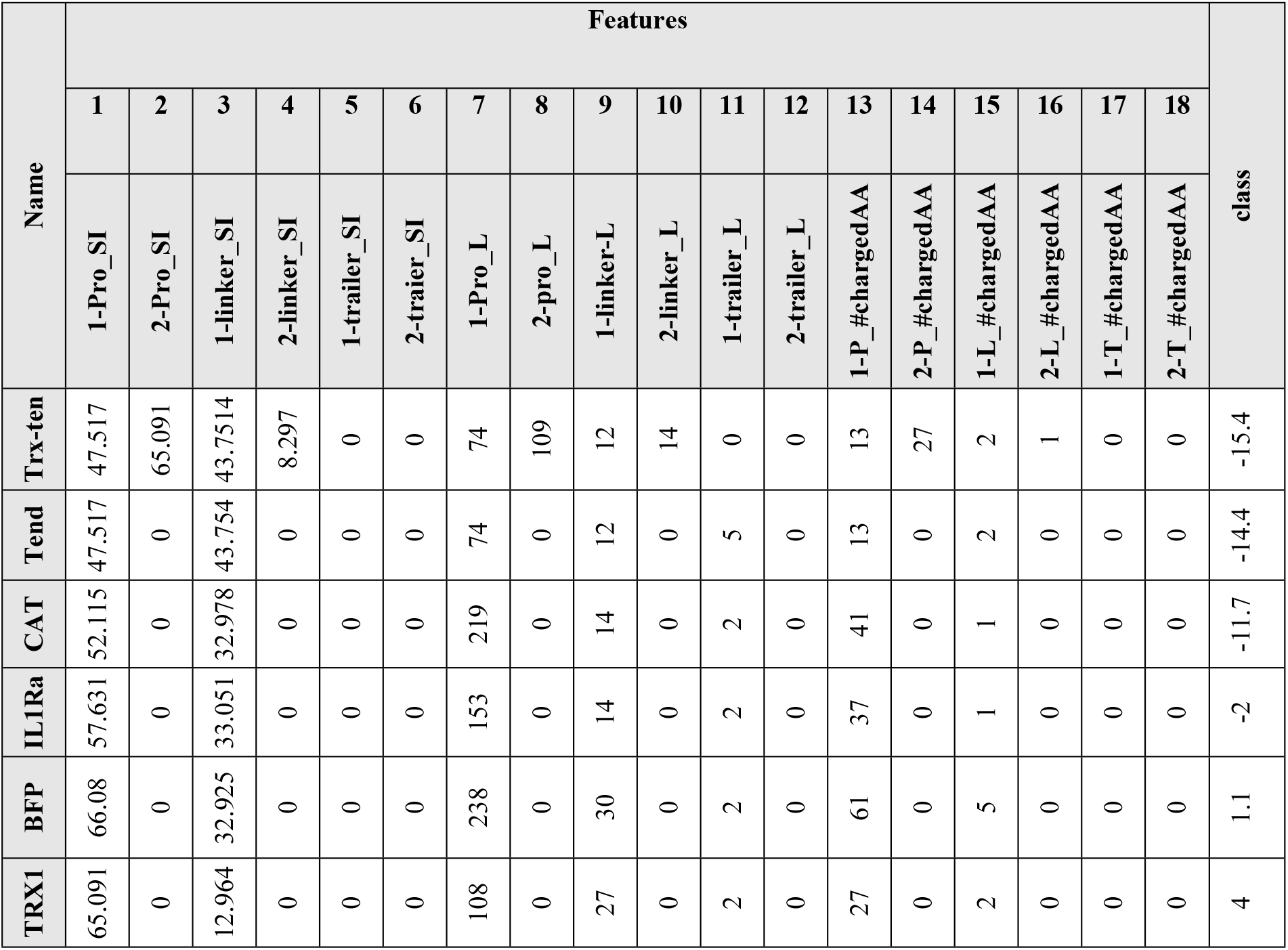
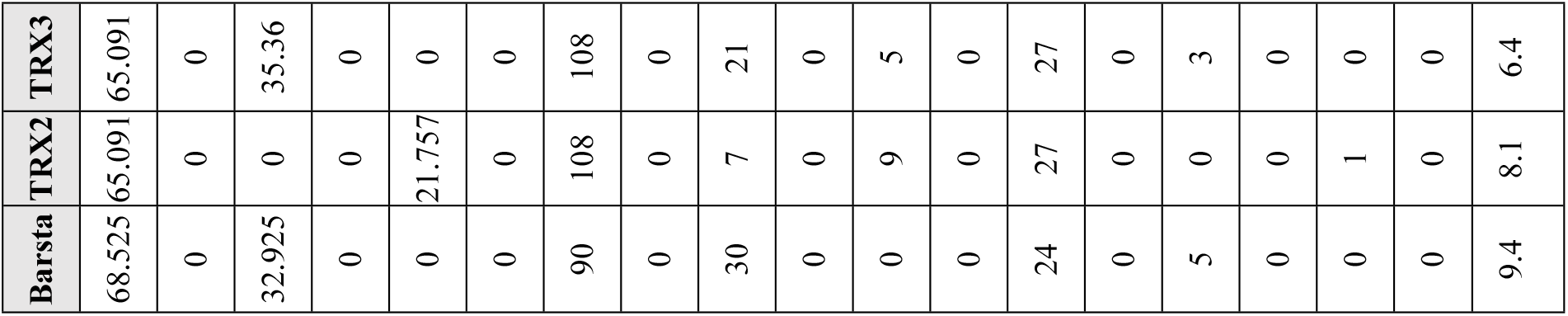
The numerically calculated values of the attributes.

### Feature selection

The feature selection paths (i.e., Select attributes option > attribute evaluator > Choose tab > WrapperSubsetEval > Classifier > function > LinearRegression) were followed in the Weka software to evaluate the attributes. On account of the experimental data, the 9-fold cross-validation was selected in the attribute mode section. After clicking on the start button, the least significant features having less than 50% of the number of folds were eliminated from the pre-process toolbar in the software. Figure 1 depicts the best effective attributes (i.e., over 50% of the number of folds).

**Fig. 1.**
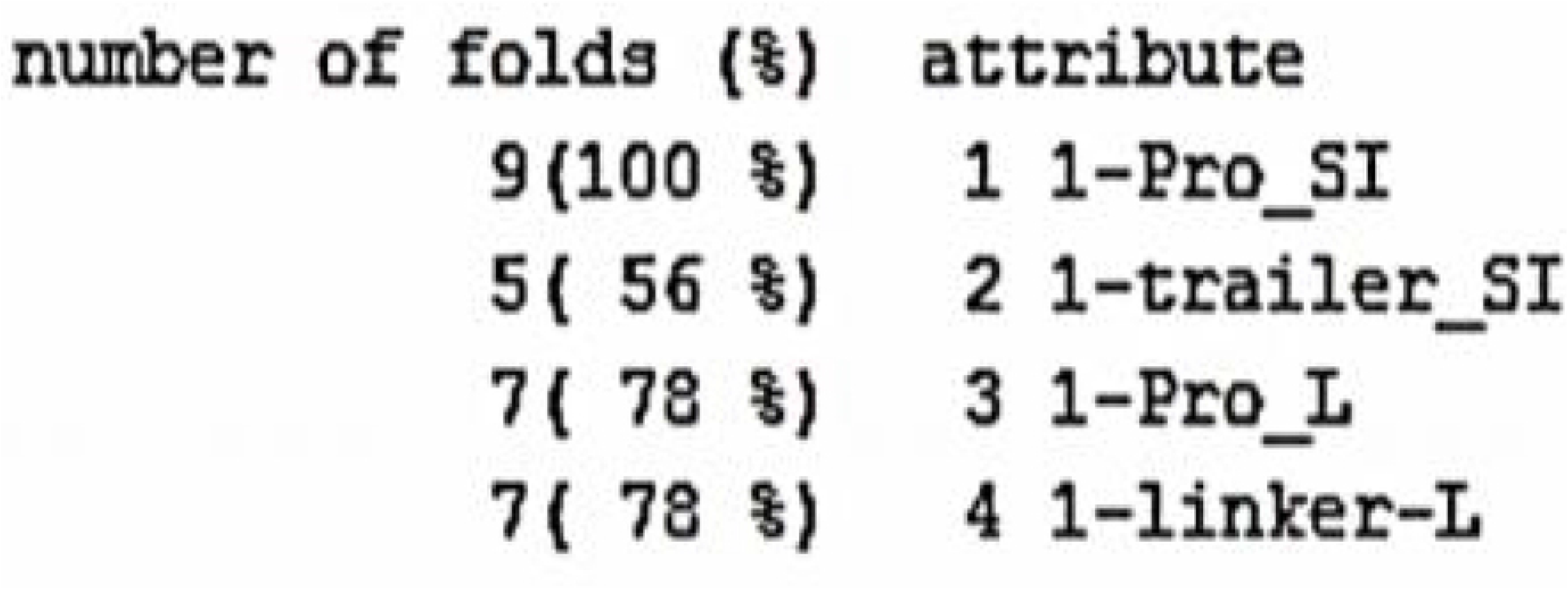
The best-selected attributes above 50% in the feature selection section of the Weka software listed at the bottom of the “number of folds (%)” column.

### Classifier

The numerical classifier selection was followed by clicking on the “classify toolbar.” The “All function” option was selected in the “choose Tab.” Then, the 9-fold cross-validation was then selected in the Weka software, and the start button was employed. Finally, the linear regression function showed the highest correlation coefficients. The result of this classifier was automatically calculated, as shown in Figure 2.

**Fig. 2.**
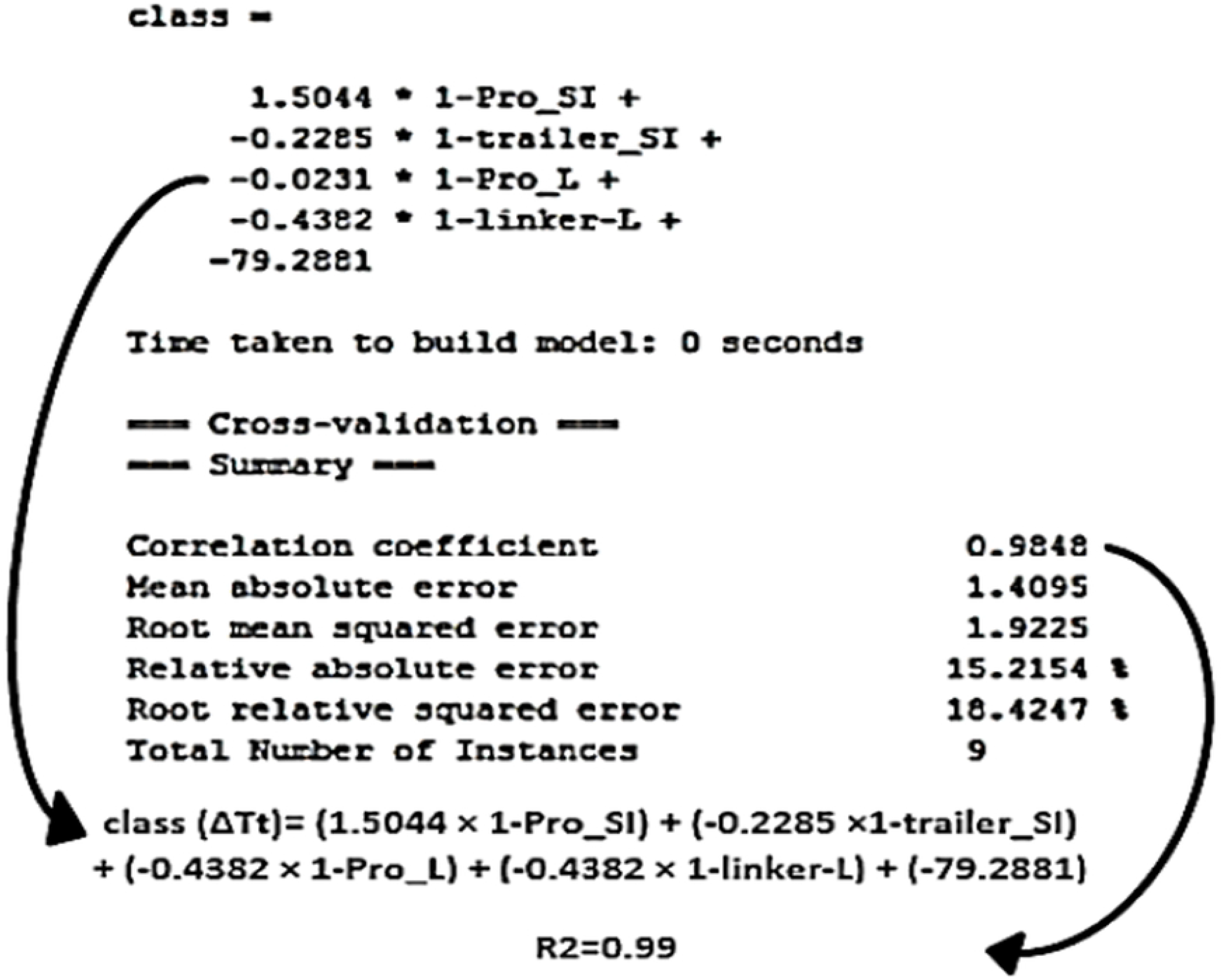
The 9-fold cross-validation result of the classify toolbar in the Weka software (This model calculated the Tt value using the 1-Pro_SI, 1-trailer_SI, 1-Pro_ L, and 1-linker_L features).

### Classifier evaluation

The numerical values of the internal controls were the previously analyzed data-set in the feature extraction step. These values indicate sample errors or bias in the model. The external control (i.e., GFP-ELP) was used for double-checking the model performance and calculated the out-of-sample errors or variances of the predictor. Table 2 represents the overall results for model assessment and the predicted and real-class values. The grey highlighted column (No. 10) demonstrates the external control in Table 2.

**TABLE 2.**
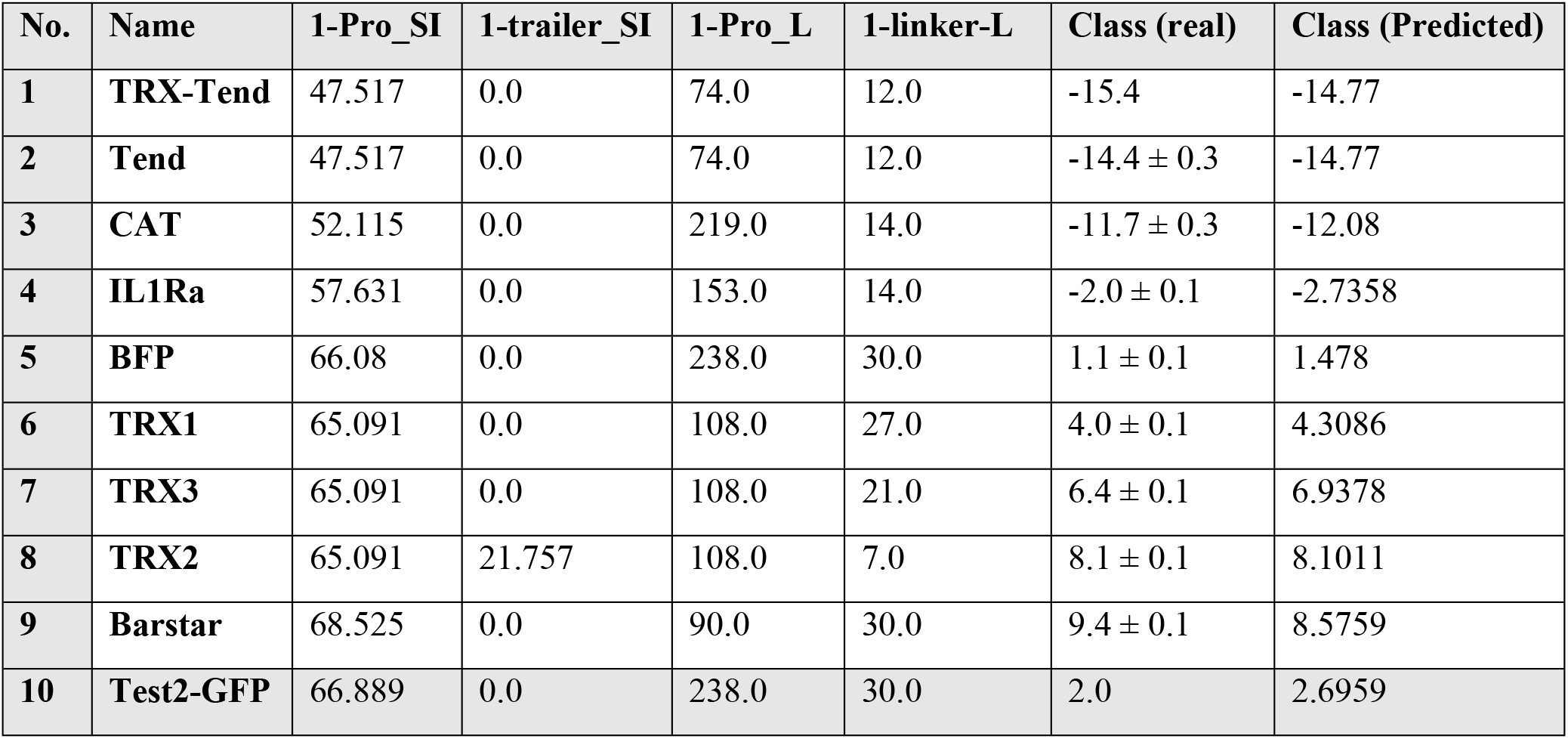
The numerically calculated results of internal and external controls. The predicted class is the ΔTt value of equation 2 measured by the predictor model represented in Figure 5. This model is briefly shown in the predictor model No. 1 shown in Table 3.

| No. | Name | 1-Pro_SI | 1-trailer_SI | 1-Pro_L | 1-linker-L | Class (real) | Class (Predicted) |
| --- | --- | --- | --- | --- | --- | --- | --- |
| 1 | TRX-Tend | 47.517 | 0.0 | 74.0 | 12.0 | -15.4 | -14.77 |
| 2 | Tend | 47.517 | 0.0 | 74.0 | 12.0 | $-14.4 \pm 0.3$ | -14.77 |
| 3 | CAT | 52.115 | 0.0 | 219.0 | 14.0 | $-11.7 \pm 0.3$ | -12.08 |
| 4 | IL1Ra | 57.631 | 0.0 | 153.0 | 14.0 | $-2.0 \pm 0.1$ | -2.7358 |
| 5 | BFP | 66.08 | 0.0 | 238.0 | 30.0 | $1.1 \pm 0.1$ | 1.478 |
| 6 | TRX1 | 65.091 | 0.0 | 108.0 | 27.0 | $4.0 \pm 0.1$ | 4.3086 |
| 7 | TRX3 | 65.091 | 0.0 | 108.0 | 21.0 | $6.4 \pm 0.1$ | 6.9378 |
| 8 | TRX2 | 65.091 | 21.757 | 108.0 | 7.0 | $8.1 \pm 0.1$ | 8.1011 |
| 9 | Barstar | 68.525 | 0.0 | 90.0 | 30.0 | $9.4 \pm 0.1$ | 8.5759 |
| 10 | Test2-GFP | 66.889 | 0.0 | 238.0 | 30.0 | 2.0 | 2.6959 |

### Reliable Analysis

Since the dataset is increased by CT-GAN method, it is more reliable to change our classifier analyzer tool. In this regard, we have used Elasticsearch engines which is a Java-based program provides by google. The Elasticsearch engine, is powered by classification-based supervised learning, uses a new type of boosted method called, boosted tree regression (24). The classification decision is based on decision tree algorithm with some special hyperparameters. In more detail, every tree helps the previous ones to improve the final iteration decision. According to our method introduced in 2.7 section, at first, we have classified our data into two types of labels, called One Fusion (OF) and Two Fusion (TF). The Figure 3 illustrates Receiver Operating Characteristic (ROC) curve evaluated by boosted Random Forest (RF) method in Elasticsearch. We have inferred that in our reliable dataset, since of having overall accuracy score of 76% in classification, OF class are more predictable in compare of the other rival which it has 64% accuracy. For more detail, the Table 3 offers an overview of the classification analysis performance, especially the normalized confusion matrix. Examining this table reveals how the categorization process mistaken the various classes when making its predictions.

**TABLE 3.** Confusion matrix for entire dataset analysis.

| Predicted Class | Actual Class |  |
| --- | --- | --- |
|  | One Fusion | Two Fusion |
| One Fusion | 76% | 24% |
| Two Fusion | 36% | 64% |

**Fig. 3.**
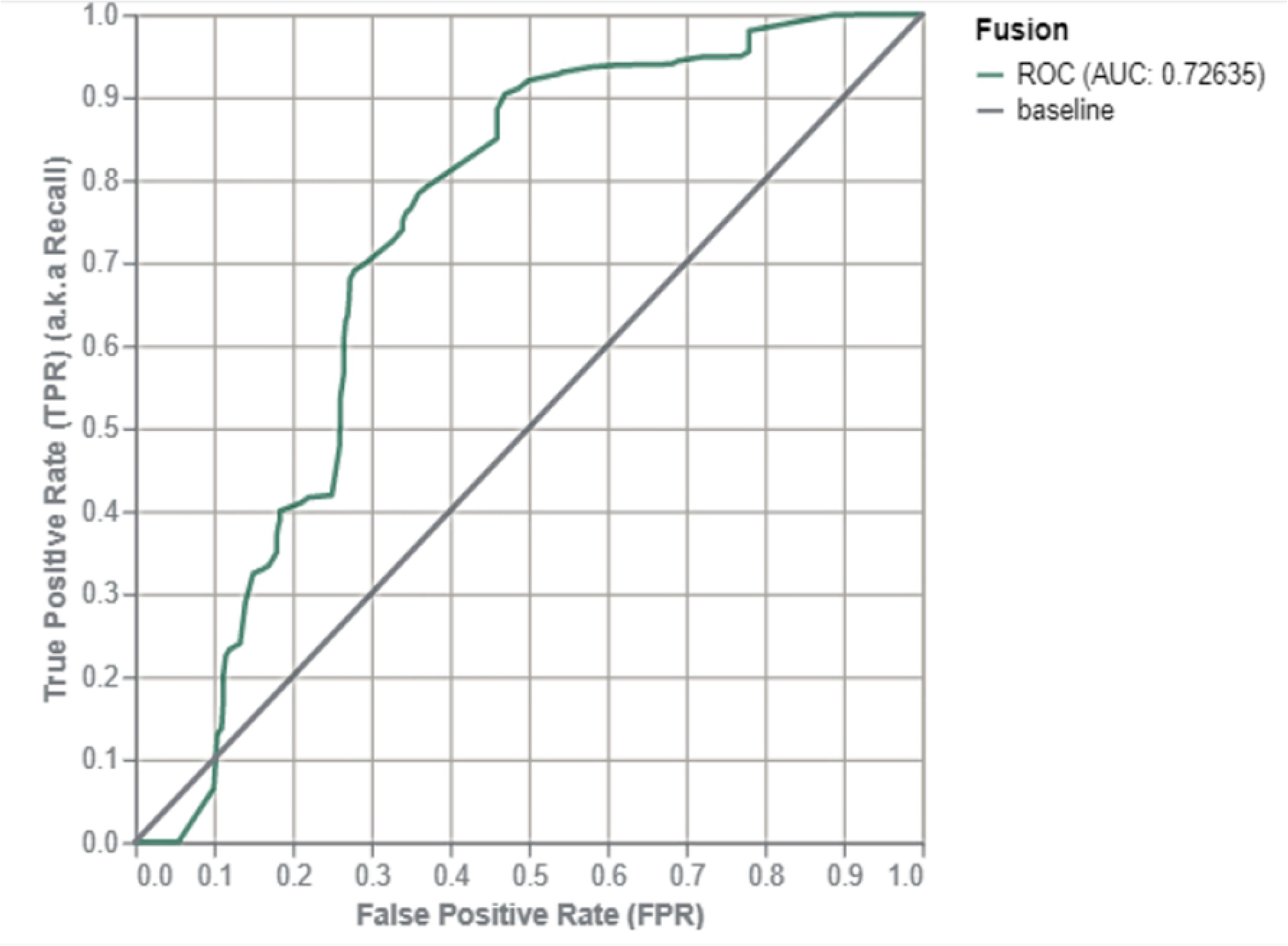
Variation of false positive rate over true positive rate.

Since of our reliable dataset, evaluating the equation achieved by No.1 Table 3 is not without merit. The Figure 4 shows the top feature importance values for regression analysis, evaluated by Elasticsearch. This result shows that in large number of proteins, 1-Pro_SI feature has the most impact on transition values, which exactly supports our achieved equation.

**Fig. 4.**
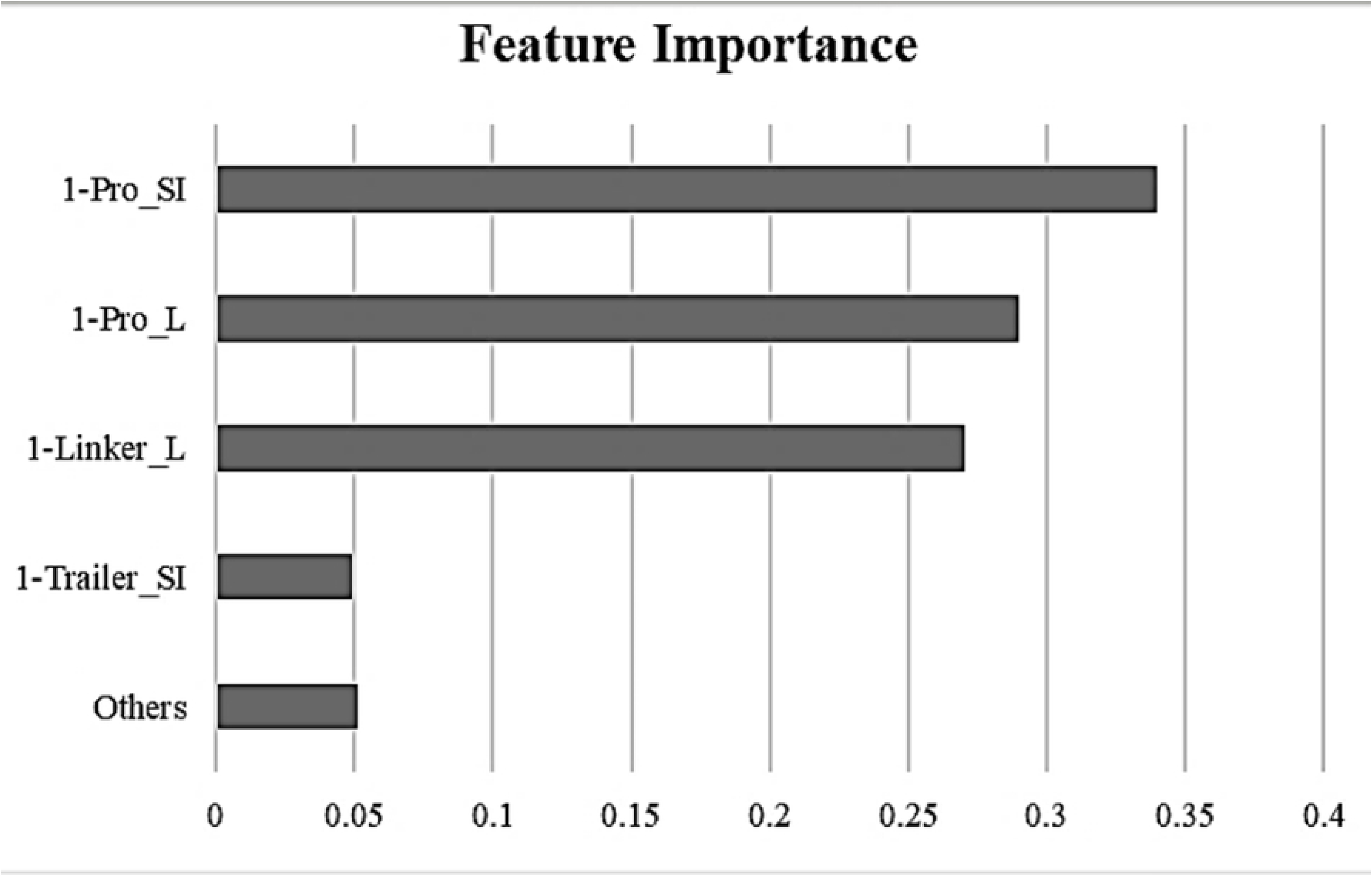
Top feature importance value for regression analysis.

## Discussion

It has been shown that the fusion of proteins, linkers, and trailers to ELP tags can affect the Tt values of the fused ELP (23). In the present study, 18 features were extracted for the protein, linker, and trailer sequences. Also, consistent with a study carried out by Christensen and colleagues (23), the charged amino acids surface index (SIc) could be used for Tt prediction of fused ELPs. After the feature analysis in the Weka software, four out of the selected 18 features showed a significant linear effect on Tt used for the classifier selection. As shown in Figure 2, the selected model could accurately predict the Tt value with R2= 0.99. Table 4 shows the predictor model for calculating the ΔTt values in our study as well as the previous report. It was obvious that the most and less effective features for the prediction of Tt were 1-Pro_SI and 1-trailer_SI, respectively, because of their coefficient differences in the model (i.e., +1.5044 and −0.2285, respectively). Moreover, as depicted in Table 5, the predicted and actual values of the Tt values were perfectly matched, and the accuracy of this model was higher than the previous model reported by Christensen and colleagues (i.e., R2=0.93) (23). Christensen et al. calculated the surface index of the BFP protein using the GFP crystal structure. They demonstrated that the BFP sequence has a higher similarity to the PDB file of GFP protein. Therefore, the PDB file for GFP was used to calculate the extinction coefficient and the SI value of BFP. Since the peptide sequences were only used to determine the SIc; the peptide sequencing information alone could predict the Tt value of fused ELPs with high accuracy. Another advantage of our model over the previous model is that it is capable of analyzing the effects of linkers and trailers (in addition to the protein features) on Tt values of the binary and ternary fused ELP constructions in the final predictor model. Therefore, in the current research, the Tt value of TRX1-ELP, TRX2-ELP, and ELP-TRX3 were predicted to be 54.30°C, 58.10°C, and 56.94°C, respectively, while the previous prediction was at the same temperature (i.e., 54.44°C) (23).

**TABLE 4.** The predictor models to calculate the numerical differences in the Tt values between the fused ELP protein and non-fused ELP. The SI values calculated by equation 1.

| NO | Predictor model | R2 | Represented in |
| --- | --- | --- | --- |
| 1 | Classifier 1 ($\Delta Tt$) = $(1.5044 \times 1\text{-Pro\_SI}) + (-0.2285 \times 1\text{-trailer\_SI}) + (-0.4382 \times 1\text{-Pro\_L}) + (-0.4382 \times 1\text{-linker-L}) - (79.2881)$ | = 0.99 | this study |
| 2 | Classifier 2 ($\Delta Tt$) = $(1.1379 \times 1\text{-Pro\_SI}) - (69.2092)$ | =0.96 | this study |
| 3 | Classifier 3 ($\Delta Tt$) = $(0.77 \times SI) - (47.4)$ | = 0.93 | the previous study by *Christensen et al* |

**TABLE 5.** The comparison of the predicted Tt values between the results of our study (i.e., Tt1 °C and Tt2 °C] and previous studies (i.e., Tt3 °C). The predicted values of Tt1, Tt2, and Tt3 show the Tt (fusion ELP) of equation 2. The predictor models No. 1, 2, or 3 in Table 4 calculated the ΔTt value of the Tt1, Tt2, and Tt3, respectively. Tt (real) is an experimental value that is previously reported (22, 23).

| No. | Name | Tt (real) °C | Tt (1) °C | Tt (2) °C | Tt (3) °C |
| --- | --- | --- | --- | --- | --- |
| 1 | TRX-Tend | 34 | 35.23 | 34.860 | 35.59 |
| 2 | Tend | 35.6 ± 0.7 | 35.23 | 34.860 | 35.59 |
| 3 | CAT | 38.3 ± 0.8 | 37.9200 | 40.093 | 40.52 |
| 4 | **IL1Ra** | 48.0 ± 0.3 | 47.2642 | 46.369 | 45.63 |
| 5 | **BFP** | 51.1 ± 0.5 | 51.478 | 55.983 | 54.17 |
| 6 | **TRX1** | 54.0 ± 0.5 | 54.3086 | 54.858 | 54.44 |
| 7 | **TRX3** | 56.4 ± 0.4 | 56.9378 | 54.858 | 54.44 |
| 8 | **TRX2** | 58.1 ± 0.8 | 58.1011 | 54.858 | 54.44 |
| 9 | **Barstar** | 59.4 ± 0.4 | 58.5759 | 58.765 | 61.24 |
| 10 | **Test2-GFP** | 52 | 52.6959 | 56.904 | 54.17 |

In the previous study, Christensen et al. used the crystal structures of proteins, along with REDUSE and PROBE software programs, to calculate the extinction coefficients and the SI values. In contrast, we do not need to use such software programs to know the peptide information, such as the PDB file, the secondary and tertiary structures of proteins. High accuracy and speed are additional advantages of this model.

In a study performed by *Chilkoti* et al., the Tt values of GFP and BFP were reported to be 52°C and 51°C, respectively (22). It is worth stating that we noted the same temperature difference after the Tt prediction in our model. With regard to the high contribution of the 1-protein_SI feature to the predictor model, another classifier (Classifier 2 in Table 4) was also introduced by only the 1-Pro_SI feature. Figure 5 illustrates another classifier to predict the Tt values that were only used by the 1-Pro_SI feature. As expected, the elimination of the linker and trailer features led to a reduction in the accuracy of the model. Nevertheless, this predictor model still had the correlation coefficient more than the previous study (i.e., 0.96 versus 0.93). These findings show that the ASA information obtained from the NetsurfP-server is able to calculate the Sic value better than REDUSE and PROBE software programs in the prediction of the Tt values. Regarding the wide application of ELPs in biological sciences, machine learning and the information obtained from the server NetsurfP can be used as a novel, simple tools for Tt prediction of the wide variety of ELPs in the future.

**Fig 5.**
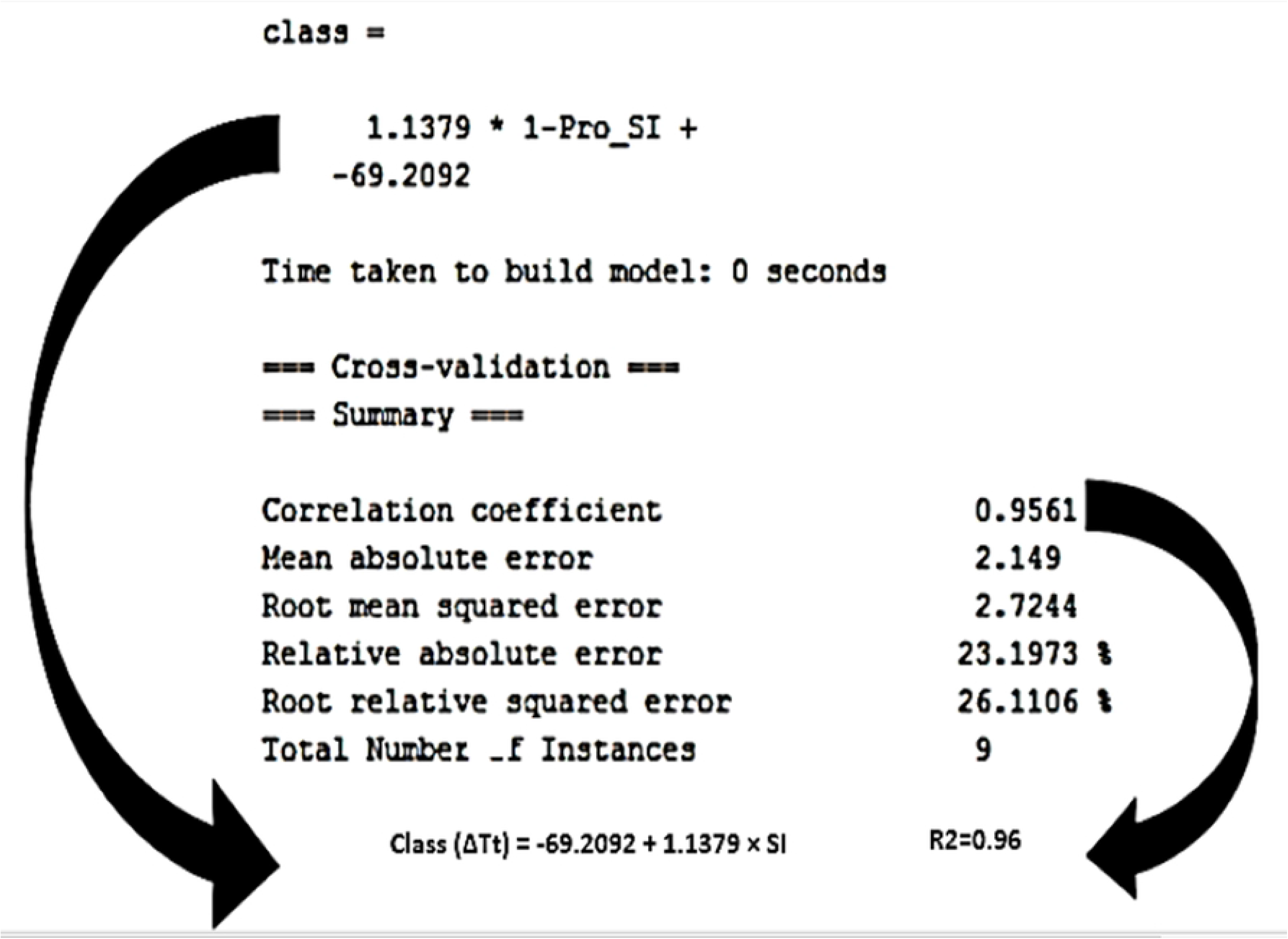
The model calculated the Tt value using the 1-Pro_SI, 1 feature.

## Materials and methods

### Structures of protein

The ELP90 protein was utilized as a fusion tag in this study. The molecular weight of ELP90 is 36kDa and has Val, Ala, or Gly (5: 2: 3) amino acids at the fourth position of the pentapeptide repeat sequence. The PDB files of the fused thioredoxin (TRX), tendamistat, chloramphenicol acetyltransferase (CAT), blue fluorescent protein (BFP), interleukin 1 receptor antagonist (IL1Ra), barstar, and green fluorescent protein (GFP) were obtained from the protein data bank with the following file names 2trx, 1ok0, 1pd5, 1bfp, 1ilr, 1a19, and 1gfl, respectively. The structures of the ELP-proteins have already been used for feature extraction/selection, cross-validation, quantitative model determination, and external control validation (22, 23). The structures of these proteins are depicted in Figure 6.

**Fig. 6.**
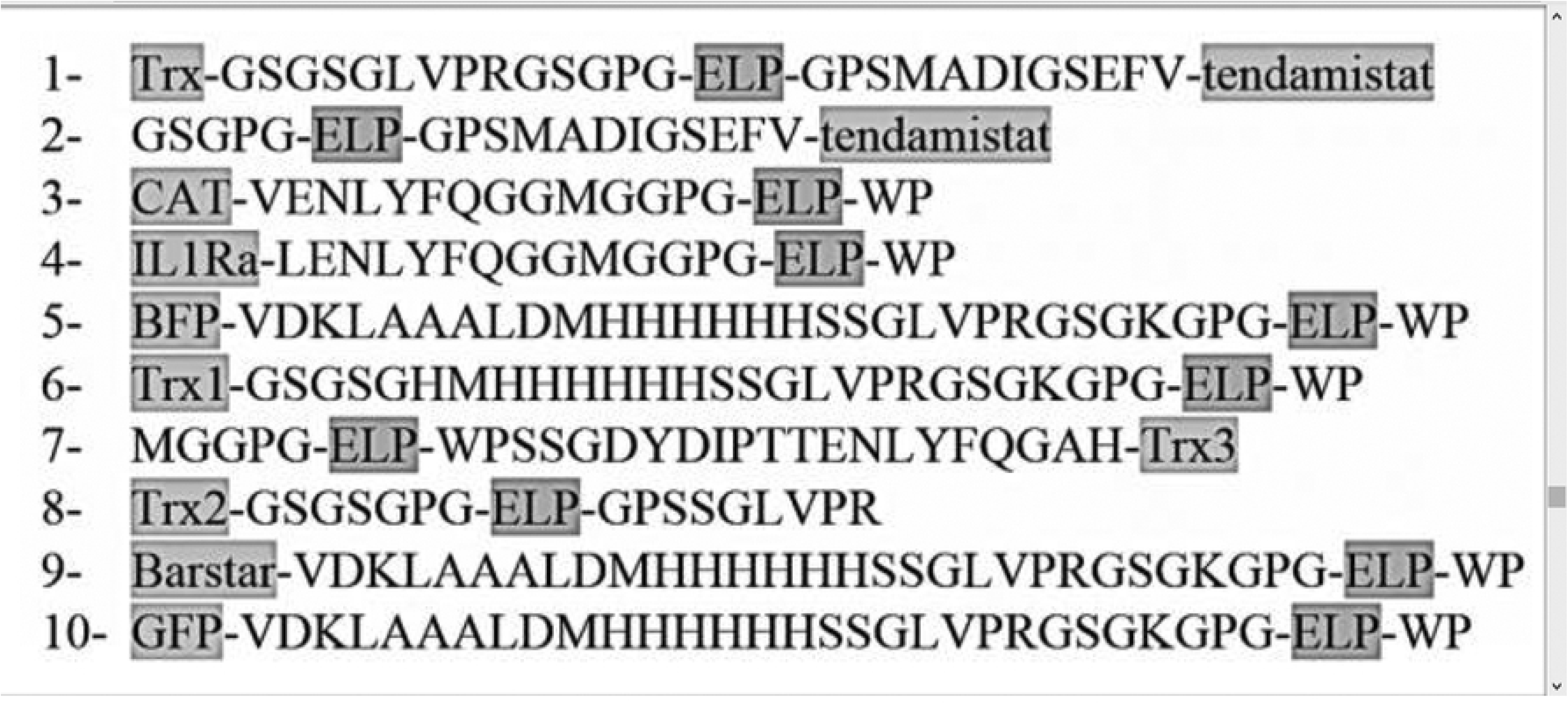
The schematic design of the fused proteins (highlighted as gray at the end of the structure), ELP (highlighted as gray at the middle of the structure), linker, and trailer (not highlighted).

### Feature extraction

Regarding the protein structures displayed in Figure 6, eighteen features were extracted. These features and the calculation methods are shown in Table 6. In Tables 6 and 1, the prefix values of 1 and 2 (i.e., 1-Pro_SIc، 2-Pro_Sic) and the other prefix values the first and second fused proteins/ linkers/trailers in the ELP-protein or protein-ELP construction. Of note, the zero numbers were used for these features with prefix 2 with the binary structure in Table 1 because there exists only one type of the protein, linker, and trailer in binary constructs. Also, the N-terminal protein, linker, and trailer were coined as prefix 2 in ternary structures. Finally, the Weka software, version 3.8.1, was employed for data mining and extraction, which is a comprehensive open-source library to perform data mining and machine learning (3, 25).

**TABLE 6.**
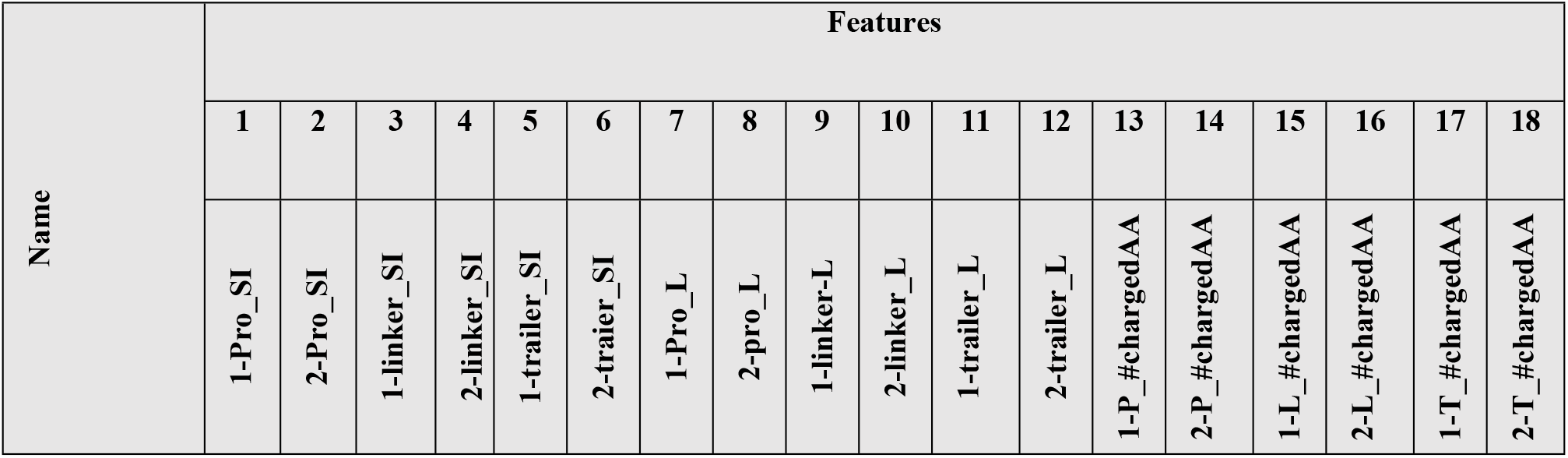
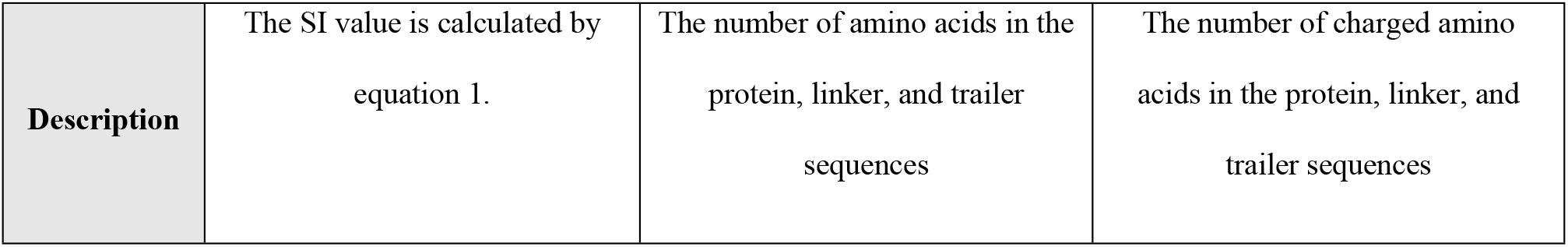
Extracted features were obtained in this experiment.

| Name | Features |  |  |  |  |  |  |  |  |  |  |  |  |  |  |  |  |  |
| --- | --- | --- | --- | --- | --- | --- | --- | --- | --- | --- | --- | --- | --- | --- | --- | --- | --- | --- |
|  | 1 | 2 | 3 | 4 | 5 | 6 | 7 | 8 | 9 | 10 | 11 | 12 | 13 | 14 | 15 | 16 | 17 | 18 |
|  | 1-Pro_SI | 2-Pro_SI | 1-linker_SI | 2-linker_SI | 1-trailer_SI | 2-trailer_SI | 1-Pro_L | 2-Pro_L | 1-linker-L | 2-linker_L | 1-trailer_L | 2-trailer_L | 1-P_#chargedAA | 2-P_#chargedAA | 1-L_#chargedAA | 2-L_#chargedAA | 1-T_#chargedAA | 2-T_#chargedAA |

|  |  |  |  |
| --- | --- | --- | --- |
| **Description** | The SI value is calculated by equation 1. | The number of amino acids in the protein, linker, and trailer sequences | The number of charged amino acids in the protein, linker, and trailer sequences |

Features 1-6 were related to surface index (SIc) in Table 6. SIc was defined as the following equation for each fused protein, linker, and trailer (23).

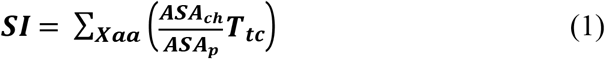

Where SI corresponds to the surface index of each fused protein, linker, and trailer. ASAch and ASAp show the accessible surface areas for a particular charged (ch) amino acid of a protein/linker/trailer (p). ASA was obtained by the free online server named NetSurfP (http://www.cbs.dtu.dk/services/NetSurfP) (26). The characteristic of the Tt value was previously measured by Urry and colleagues for any amino acid (23).

### Class definition

In the machine learning method, the class is defined as a final qualitative or quantitative output predicted. The numerical differences in the Tt values between the fused ELP protein and non-fused ELP were determined as the class in this study as represented in equation 2 (22, 23).

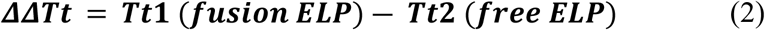

In this equation, the Tt1 and Tt2 values represent the transition temperatures for the ELP-fused protein and free ELP, respectively. Moreover, the transition temperature value of the free ELP or its free ELP construction was precisely considered the same as the value or the structure reported in the previous study (i.e., 50.0 ± 0.4 and GSGKGPG-ELP-WP).

### Feature selection

The aim of feature selection is the screening of features to find the top effective attributes in the model. Weka software has automated attribute selection facilities that are able to screen the numerical attributes by “CfsSubsetEval”, “PrincipalComponet”, “ReliefFAttributeEval” and “WrapperSubsetEval” evaluators. Considering the previous report on the linear effect of fused proteins on the Tt value of ELP, the “WrapperSubsetEval” evaluator was applied to screen the attributes. For feature selection, 18 attributes and one class were submitted to Weka software.

### Classifier

The best effective attributes selected in the previous step were applied in the classifier selection step. Except for the best effective attributes, the other attributes were removed from the pre-process option. Then, the nine-fold cross-validation and linear regression were chosen in the classification mode. Finally, the best quantitative model was obtained from the classification mode in the Weka software (3, 25).

### External controls

Following the classifier selection, numerical values of the class were predicted for the internal (i.e., Trx-ELP-ten, ELP-ten, CAT-ELP, IL1Ra-ELP, BFP-ELP, TRX1-ELP, TRX2-ELP, ELP-TRX3, and Barstar-ELP) and external controls (i.e., GFP-ELP). These predicted class values were compared with the previously calculated actual class values and then used for model performance assessment (17, 22, 23).

### Resampling and evaluation

Due to more reliability of our experimental dataset, we have extended our dataset by state-of-the-art artificial intelligence method. Generative Adversarial Networks (GAN) have been used for many years for creating fake images, data and etc. (27). In this regard, since our dataset are included by both numerical and Boolean values, we have used a novel GAN method, namely CT-GAN (28) for increasing the reliability of our dataset. By this method, we have increased 90 fake values in order to have a clean and complete dataset (Refer to the resample-2.csv in the Supplementary File). The Figure 7 illustrates our proposed methodology for creating a reliable dataset.

**Fig. 7.**
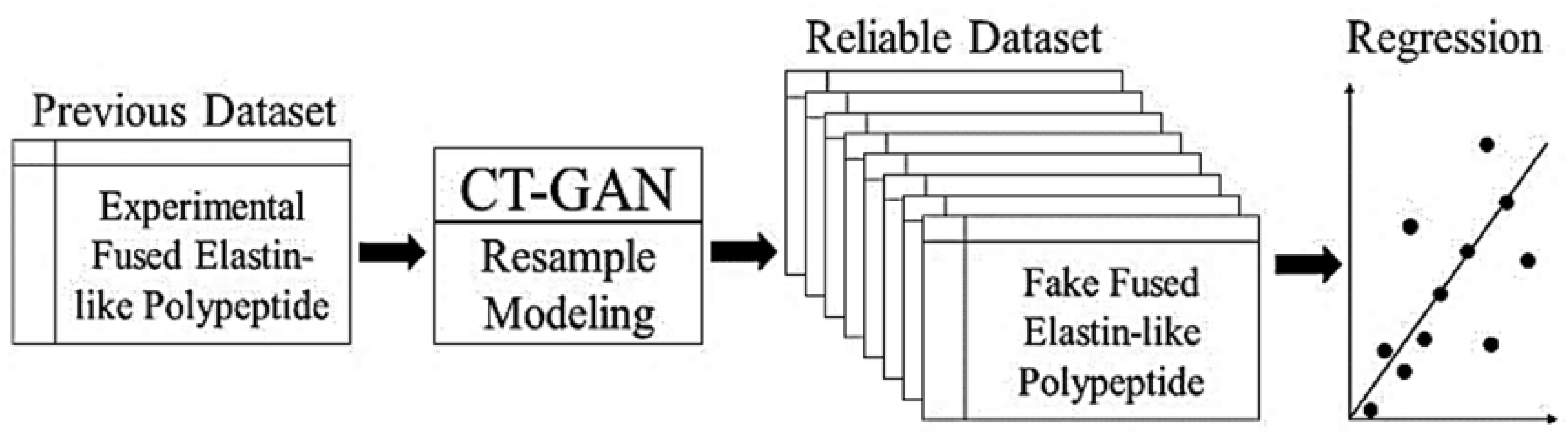
The processing of creating reliable dataset by CT-GAN resampling method.

## Supporting information

The detailed information of ASA, SI calculations and resample data is shown in the S1 Appendix. Appendix (.pdf and .csv).

## Acknowledgments

The authors are grateful to Shahed University for support of this work. The authors are also grateful to Cathal O’Seanain for re-reading and language/editing assistance.

## References

1. Saxena R, Nanjan MJ. Elastin-like polypeptides and their applications in anticancer drug delivery systems: a review. Drug Deliv. 2015;22(2):156–67.

2. Noguera-Salvà MA, Guardiola-Serrano F, Martin ML, Marcilla-Etxenike A, Bergo MO, Busquets X, et al. Role of the C-terminal basic amino acids and the lipid anchor of the Gγ2 protein in membrane interactions and cell localization. Biochimica et Biophysica Acta (BBA) - Biomembranes. 2017;1859(9, Part B):1536–47.

3. Silvius JR. Lipid Modifications of Intracellular Signal-Transducing Proteins. Journal of Liposome Research. 1999;9(1):1–19.

4. Pedersen TB, Kaasgaard T, Jensen MØ, Frokjaer S, Mouritsen OG, Jørgensen K. Phase behavior and nanoscale structure of phospholipid membranes incorporated with acylated C14-peptides. Biophysical journal. 2005;89(4):2494–503.

5. Aladini F, Araman C, Becker CFW. Chemical synthesis and characterization of elastin-like polypeptides (ELPs) with variable guest residues. Journal of Peptide Science. 2016;22(5):334–42.

6. Rincón AC, Molina-Martinez IT, de las Heras B, Alonso M, Baílez C, Rodríguez-Cabello JC, et al. Biocompatibility of elastin-like polymer poly(VPAVG) microparticles: in vitro and in vivo studies. Journal of Biomedical Materials Research Part A. 2006;78A(2):343–51.

7. Urry DW, Pattanaik A, Jie X, Cooper Woods T, McPherson DT, Parker TM. Elastic protein-based polymers in soft tissue augmentation and generation. Journal of Biomaterials Science, Polymer Edition. 1998;9(10):1015–48.

8. Kenworthy A. Peering inside lipid rafts and caveolae. Trends Biochem Sci. 2002;27(9):435–7.

9. Meyer DE, Chilkoti A. Quantification of the Effects of Chain Length and Concentration on the Thermal Behavior of Elastin-like Polypeptides. Biomacromolecules. 2004;5(3):846–51.

10. Urry DW. Free energy transduction in polypeptides and proteins based on inverse temperature transitions. Prog Biophys Mol Biol. 1992;57(1):23–57.

11. Urry DW. Physical Chemistry of Biological Free Energy Transduction As Demonstrated by Elastic Protein-Based Polymers. The Journal of Physical Chemistry B. 1997;101(51):11007–28.

12. Li B, Alonso DO, Daggett V. The molecular basis for the inverse temperature transition of elastin. J Mol Biol. 2001;305(3):581–92.

13. Li B, Alonso DOV, Bennion BJ, Daggett V. Hydrophobic Hydration Is an Important Source of Elasticity in Elastin-Based Biopolymers. Journal of the American Chemical Society. 2001;123(48):11991–8.

14. Li B, Daggett V. The molecular basis of the temperature- and pH-induced conformational transitions in elastin-based peptides. Biopolymers. 2003;68(1):121–9.

15. Toonkool P, Jensen SA, Maxwell AL, Weiss AS. Hydrophobic Domains of Human Tropoelastin Interact in a Context-dependent Manner*. Journal of Biological Chemistry. 2001;276(48):44575–80.

16. Meyer DE, Chilkoti A. Genetically encoded synthesis of protein-based polymers with precisely specified molecular weight and sequence by recursive directional ligation: examples from the elastin-like polypeptide system. Biomacromolecules. 2002;3(2):357–67.

17. Trabbic-Carlson K, Liu L, Kim B, Chilkoti A. Expression and purification of recombinant proteins from Escherichia coli: Comparison of an elastin-like polypeptide fusion with an oligohistidine fusion. Protein Sci. 2004;13(12):3274–84.

18. Hassouneh W, Christensen T, Chilkoti A. Elastin-like polypeptides as a purification tag for recombinant proteins. Curr Protoc Protein Sci. 2010;Chapter 6:Unit 6.11.

19. Huang KZ, Xiong XK, Zhang CM, Lai YY, Zou CN, Zhang GY, et al. Enhancement predicting accuracy for elastin-like polypeptides temperature transition by back propagation neural network. Protein Pept Lett. 2014;21(10):1065–72.

20. McDaniel JR, Radford DC, Chilkoti A. A Unified Model for De Novo Design of Elastin-like Polypeptides with Tunable Inverse Transition Temperatures. Biomacromolecules. 2013;14(8):2866–72.

21. Meyer DE, Chilkoti A. Purification of recombinant proteins by fusion with thermally-responsive polypeptides. Nat Biotechnol. 1999;17(11):1112–5.

22. Trabbic-Carlson K, Meyer DE, Liu L, Piervincenzi R, Nath N, LaBean T, et al. Effect of protein fusion on the transition temperature of an environmentally responsive elastin-like polypeptide: a role for surface hydrophobicity? Protein Eng Des Sel. 2004;17(1):57–66.

23. Christensen T, Hassouneh W, Trabbic-Carlson K, Chilkoti A. Predicting transition temperatures of elastin-like polypeptide fusion proteins. Biomacromolecules. 2013;14(5):1514–9.

24. Lu Z, Chen C, Xin J, Yu Z. On the Auto-Tuning of Elastic-search based on Machine Learning. 2020 International Conference on Control, Robotics and Intelligent System. 2020.

25. Yeboah A, Cohen RI, Rabolli C, Yarmush ML, Berthiaume F. Elastin-like polypeptides: A strategic fusion partner for biologics. Biotechnol Bioeng. 2016;113(8):1617–27.

26. Petersen B, Petersen TN, Andersen P, Nielsen M, Lundegaard C. A generic method for assignment of reliability scores applied to solvent accessibility predictions. BMC Struct Biol. 2009;9:51.

27. Creswell A, White T, Dumoulin V, Arulkumaran K, Sengupta B, Bharath AA. Generative Adversarial Networks: An Overview. IEEE Signal Processing Magazine. 2018;35(1):53–65.

28. Xu L, Skoularidou M, Cuesta-Infante A, Veeramachaneni K, editors. Modeling Tabular data using Conditional GAN. NeurIPS; 2019.

